# In silico model suggests that interdigitation promotes robust activation of atrial cells by pacemaker cells

**DOI:** 10.1101/2024.05.15.594103

**Authors:** Martijn A. de Jong, Roeland M.H. Merks

## Abstract

The heartbeat is initiated by electrical pulses generated by a specialized patch of cells called the sinoatrial node (SAN), located on top of the right upper chamber, and then passed on to the atrium. Cardiac arrhythmias may arise if these electrical pulses fail to propagate toward the atrial cells. This computational modeling study asks how the morphology of the interface between sinoatrial (pacemaker) cells and atrial cells can influence the robustness of pulse propagation. Due to its strong negative potential, the atrium may suppress the pacemaker activity of the SAN if the electrical coupling between atrial cells is too strong. If the electrical coupling is too weak, however, the pacemaker cells cannot activate the atrial cells due to a source-sink mismatch. The SAN and atrium are connected through interdigitating structures, which are believed to contribute to the robustness of action potentials and potentially solve this trade-off. Here we investigate this interdigitation hypothesis using a hybrid model, integrating the cellular Potts model (CPM) for cellular morphology and partial-differential equations-based electrophysiological models for pulse propagation. Systematic examination of interdigitation patterns revealed that a symmetric geometry with medium-sized protrusions can prevent exit blocks. We conclude that interdigitation of SAN cells and atrial cells can promote robust propagation of action potentials toward the atrial tissue but only if the protrusions are of sufficient size and synchronicity of the action potential wave is maintained due to symmetry. This study not only highlights essential design principles for *in vitro* models of cardiac arrhythmias, but also provides insights into the occurrence of exit blocks *in vivo*.

**Author summary:** Our hearts beat automatically and robustly. This autonomous heartbeat is initiated by electrical pulses generated by a specialized patch of cells called the sinoatrial node, located on top of the right upper chamber. These pulses can be interpreted as electrical signals that allow the heart muscle to contract. The heart muscle cells surrounding the sinoatrial node tend to hinder this spontaneous activation because of a mismatch in electrical properties. Therefore, the pacemaker cells must be sufficiently electrically insulated from their surroundings. However, full insulation of the pacemaker cells would hinder propagation of the activation pulse toward the rest of the heart. A common hypothesis is that the sinoatrial node is fully insulated, except for some specialized pathways. We have studied the arrangement of different cell types within these pathways with the central question: how should the sinoatrial node and atrium be connected to ensure robust propagation of the electrical pulse? We implemented a computational model inspired by *in vitro* experimental setups and found several relevant mechanisms. For example, we found that a folding-finger-like structure between the cell types can dramatically improve the robustness of action potentials propagating in such a tissue, provided that the folds do not become too small. This study may help improve design of *in vitro* models of sinoatrial node diseases.

## Introduction

The most important biological pacemaker of vertebrate hearts is located at the sinoatrial node (SAN). The pacemaker cells within this node rhythmically produce action potentials, that propagate into the surrounding right atrium (RA) tissue, thus driving contraction of the atrium. Failed propagation of action potentials from the SAN into the right atrium is known as an exit block and can lead to tachy-brady syndrome or SAN arrest [1–3]. Exit blocks would occur if the electrical coupling between the SAN and the RA would be too weak [4]. A further cause of arrhythmias is the fact that atrial cells have a lower resting potential than SAN cells [5] such that, in the absence of any insulation, the hyperpolarizing influence from the surrounding RA would suppress [6, 7], or even fully inhibit [8] the spontaneous pacing of the SAN. Such mismatch of resting potentials between the SAN and the RA is also known as ‘source-sink’ mismatch (reviewed in Ref. [9]).

Several mechanisms have been hypothesized for how the SAN can reduce the source-sink mismatch and thus prevent exit blocks. The simplest hypothesis is the gradient model as first suggested by Joyner and Van Capelle [8]. Their mathematical model suggested that weak coupling between center SAN cells, combined with stronger coupling in the periphery in the SAN, sufficed to shield the SAN center from hyperpolarization, while simultaneously allowing the SAN to drive the surrounding atrial cells. In many mammals, including humans, the SAN center contains fewer and smaller gap junctions than the periphery, suggesting that cells in the SAN periphery are indeed electrically coupled more strongly than cells in the SAN center (reviewed in Ref. [10]). This observation is consistent with the gradient model. Furthermore, the SAN is electrically insulated from the RA, thus protecting the SAN from the negative resting potential in the RA. Action potentials move from the SAN to the RA through discrete channels of connecting cells called sinoatrial conduction pathways (SACPs) [1, 2, 11, 12], also known as SAN exit pathways (SEPs) [13–15]. In a simulation study it was predicted that a gradient of cellular connectivity combined with a mosaic-like gradient of cell type prevalence within the SACPs helps prevent exit blocks. [15]. Altogether, there is clear evidence for connexin expression gradients between the SAN and the RA, and computational modeling suggests that the resulting electrical coupling gradients alleviate the risk for exit blocks.

Apart from connexin expression gradients, additional mechanisms may contribute to a gradual transition between the SAN and the RA. One hypothesis is the existence of transitional cells between the SAN and the RA [16, 17], which express ion channel proteins typical for SAN cells alongside ion channel proteins typical for atrial cells [18]. Such transitional cells could provide a coupling gradient similar to those provided by connexin gradients. Transitional cells have been observed in rabbit [19–21], dog [16] and human [18, 22], but not in guinea pig [23] or mouse [7].

Another hypothesis, not mutually exclusive to transitional cell types, was proposed by ten Velde et al. [23] who observed interdigitated interfaces between the SAN cells and atrial cells in guinea pig. Such interdigitation of SAN and atrial cells has also been observed in rabbit [21], mouse [10, 24] and human [4, 11, 25–27], and could result in an effective coupling gradient. Winslow and Varghese [28] tested the interdigitation hypothesis in an *in silico* study. They compared a round SAN with atrial protrusions and a SAN without protrusions. The SAN with protrusions enabled propagation from the SAN into the atrial cells when the atrial cells were strongly coupled while conduction failed without interdigitation. Winslow and Varghese concluded that this improved conduction was possible because tips of these atrial cells could activated effectively without hyperpolarizing the SAN. Huang and Cui [29] tested interdigtation in a quasi-1D setting. The heart was modeled as a strand of cells. Extra connections between the atrial tissue and SAN were added to model atrial protrusions. They also found that action potentials propagated more effectively if these atrial strands were added, although the SAN was also suppressed more. They concluded that invading strands of atrial cells deep into the SAN can improve the robustness of action potential propagation the most. This manuscript aims to further analyze the role of interdigitation of SAN and atrium cells in preventing exit blocks. We studied the morphology of interdigitation by studying a wide variety of protrusions shapes in 2D simulations to identify on which shapes action potentials propagate most robustly. Furthermore, the effect of interdigitation was isolated from other effects that potentially influence to robustness of action potential propagation. Based on these insights, we aim to propose design principles for *in vitro* models of SAN and atrium cells, in particular suggesting shapes providing optimal electrical coupling between SAN and atrial cells.

To investigate the effect of interdigitation in the coupling between sinoatrial and atrial cells, we implemented a two-dimensional mathematical model. In this model, we study propagation of depolarization waves initiated in SAN cells through an interdigitated tissue interface, toward a simulated atrial tissue. The mathematical model combines a cellular level representation of the SAN, atrium and SAN-atrium interface with electrophysiological models that are resolved at subcellular level. The model set-up is based on typical *in vitro* set-ups [30]. In such set-ups, a tissue of SAN cells is coupled to a tissue of atrial cells via a narrow channel called the isthmus, similar to the simulation set-up in [12]. This configuration was chosen to mimic the connection of the SAN to the right atrium by SACPs in a simplified way. To focus on the effect of interdigitation, we have limited the model to two cell types, thus ignoring the hypothetical transitional cell types. In the remainder of this paper, we start by studying wave propagation on irregular boundaries resembling SAN-atrium interdigitation, which is generated using a Cellular Potts model (CPM). Surprisingly, initial simulations suggest that at such irregularly interdigitated SAN-atrium interfaces, exit blocks occur more frequently than on regular boundaries. Therefore, we next characterized three main properties of irregular tissue boundaries generated in the Cellular Potts model. Cell mixing at tissue interface led to: (1) increased boundary length, (2) interdigitation, and (3) non-straight boundaries leading to asynchronous arrival of action potentials to the atrial tissue. We study these three effects in isolation from on another, and find that long boundaries with symmetric, medium-sized protrusions lead to the most robust propagation of action potentials.

## Materials and methods

### Conceptual model

The sinoatrial node is fully insulated from the right atrium, except for several SACPs. We have modeled one SACP as a narrow tissue that couples a part of the sinoatrial node with a part of the right atrium, see (Fig 1A). The cardiac tissue surrounding the the sinoatrial node is inhomogeneous. The most important cell types in and around the sinoatrial node are specialized pacemaker cells, atrial cells, and fibroblasts [11].

**Fig 1.**
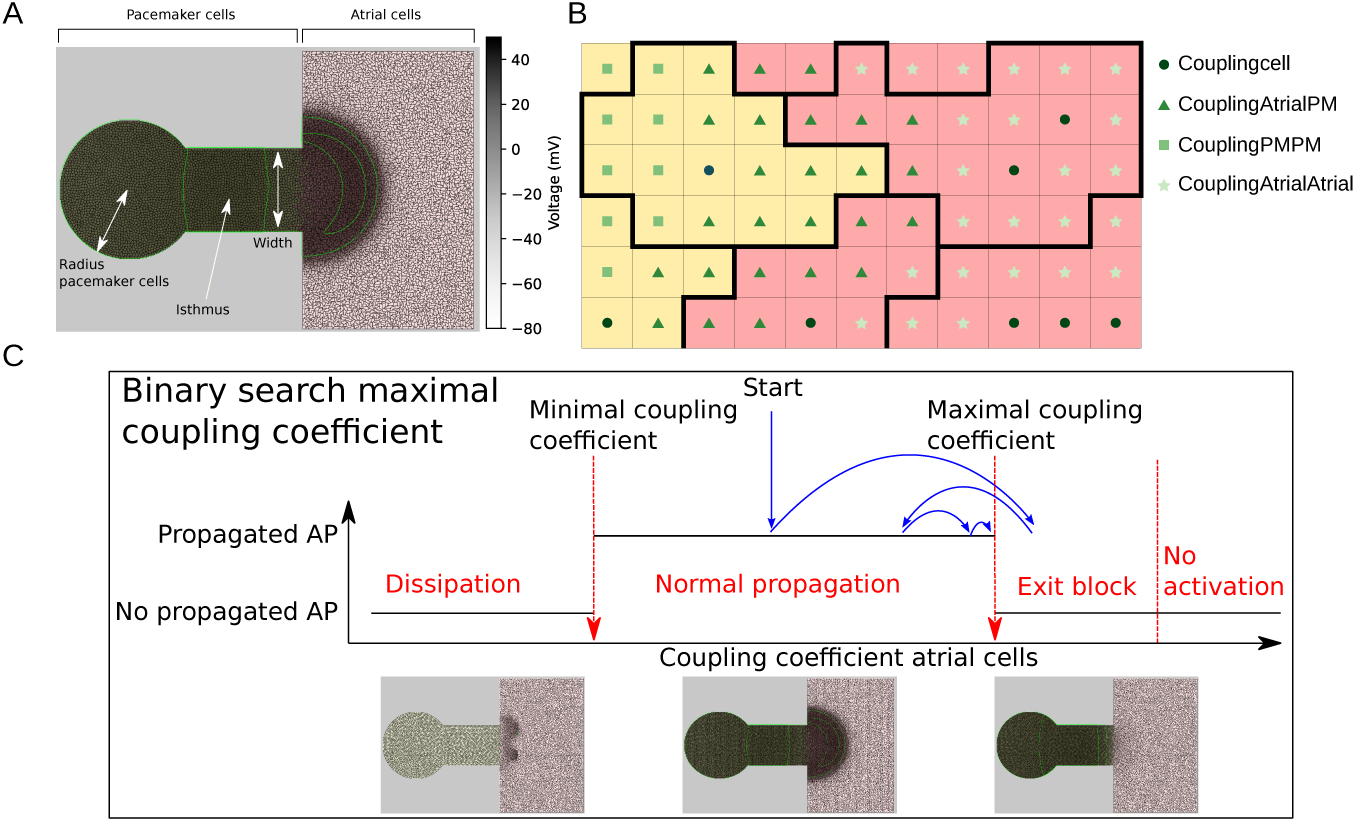
Methods used for simulations and analysis. A: Configuration of the simulations. Pacemaker cells are located within a circle on the left and in the isthmus, indicated by yellow. Atrial cells are located in the rectangle on the right, indicated by red. The shades indicate the voltage, with green lines indicating locations of equal voltage. B: A sketch of the four different coupling coefficients in configurations. Couplingcell is assigned to pixels in a cell’s interior, couplingAtrialPM to pixels at the boundary of the two cell types, couplingPMPM to pixels at the boundary between two pacemaker cells and couplingAtrialAtrial to pixels at the boundary between two atrial cells. C: A sketch of the binary to find the maximal coupling coefficient. If the coupling coefficient is too low, the action potential wave dissipates. On the other hand, if the coupling coefficient is too high, there is an exit block or no activation of the pacemaker cells. The maximal coupling coefficient is the highest value such that the action potential is propagated and is found by a binary search.

Individual pacemaker cells are ovoid or round, and 5 to 10 µm in diameter [16]. Atrial cells are typically much larger and more elongated at 6-8 µm in width by 20-30 µm in length [31]. Fibroblasts, are only modeled implicitly, by limiting the connection between pacemaker cells and atrial cells to a narrow isthmus.

Pacemaker and atrial cells differ electrophysiologically from each other: Pacemaker cells activate spontaneously and action potentials travel slowly through the SAN. Atrial cells on the other hand require external activation from pacemaker cells or other atrial cells and action potentials propagate much faster through the right atrium than in the SAN. We therefore used different electrophysiological models for pacemaker cells and atrial cells. Pacemaker cells were modeled with the autocycling Fabbri-Severi model [32] and atrial cells were modeled with the atrial cells model by Maleckar et al. [33]. Furthermore, we modeled inhomogeneous expression of connexins and its effects on conduction velocity explicitly by spatially inhomogeneous electrical coupling between neighboring pixels.

Using this baseline model, we asked how the shape of the interface between pacemaker and atrial cells could affect propagation of action potentials from the SAN to the atrium. The interfaces were generated with two methods. We studied boundaries that evolved from random cell motility in (Figs 2 and 3) and fixed the boundary geometry in (Figs 4 and 5). The robustness of action potentials at these interfaces was studied both locally with a measure called ‘safety factor’, and globally by finding the largest sink that could still be activated.

**Fig 2.**
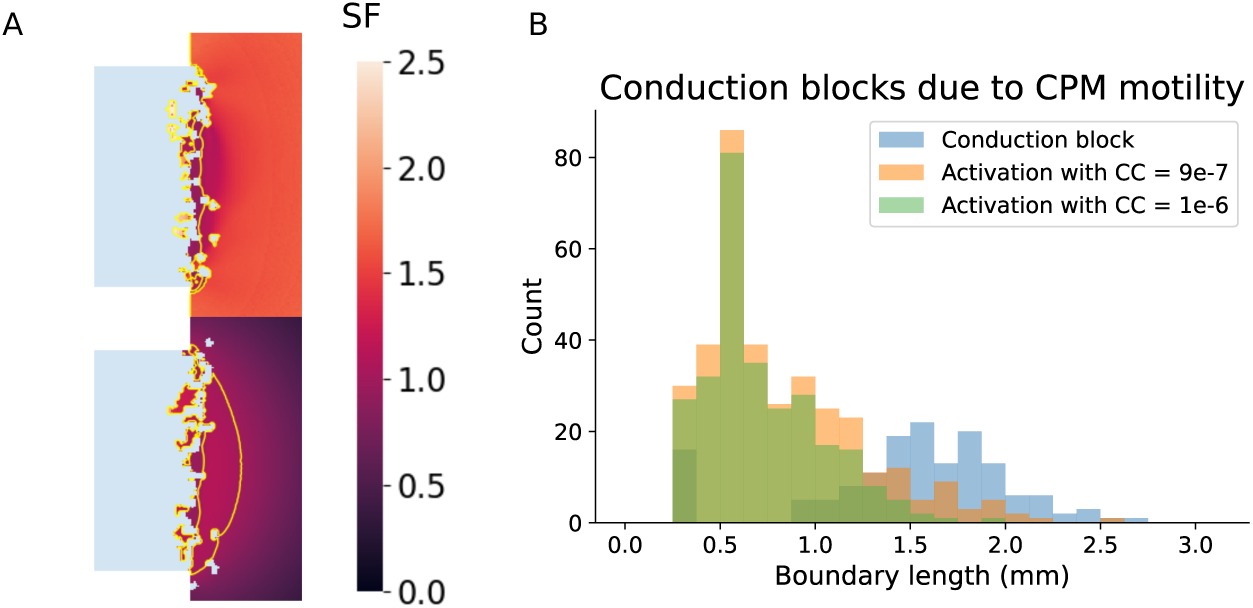
Reduced propagation of action potentials over diffuse pacemaker-atrium interfaces. A: Example of two safety factor plots for successful and failed propagation. The yellow lines in the safety factor plots show the locations where SF = 1 is achieved. B: Number of action potentials that successfully propagated on a simulated diffuse pacemaker-atrium boundaries as a function of boundary length.

**Fig 3.**
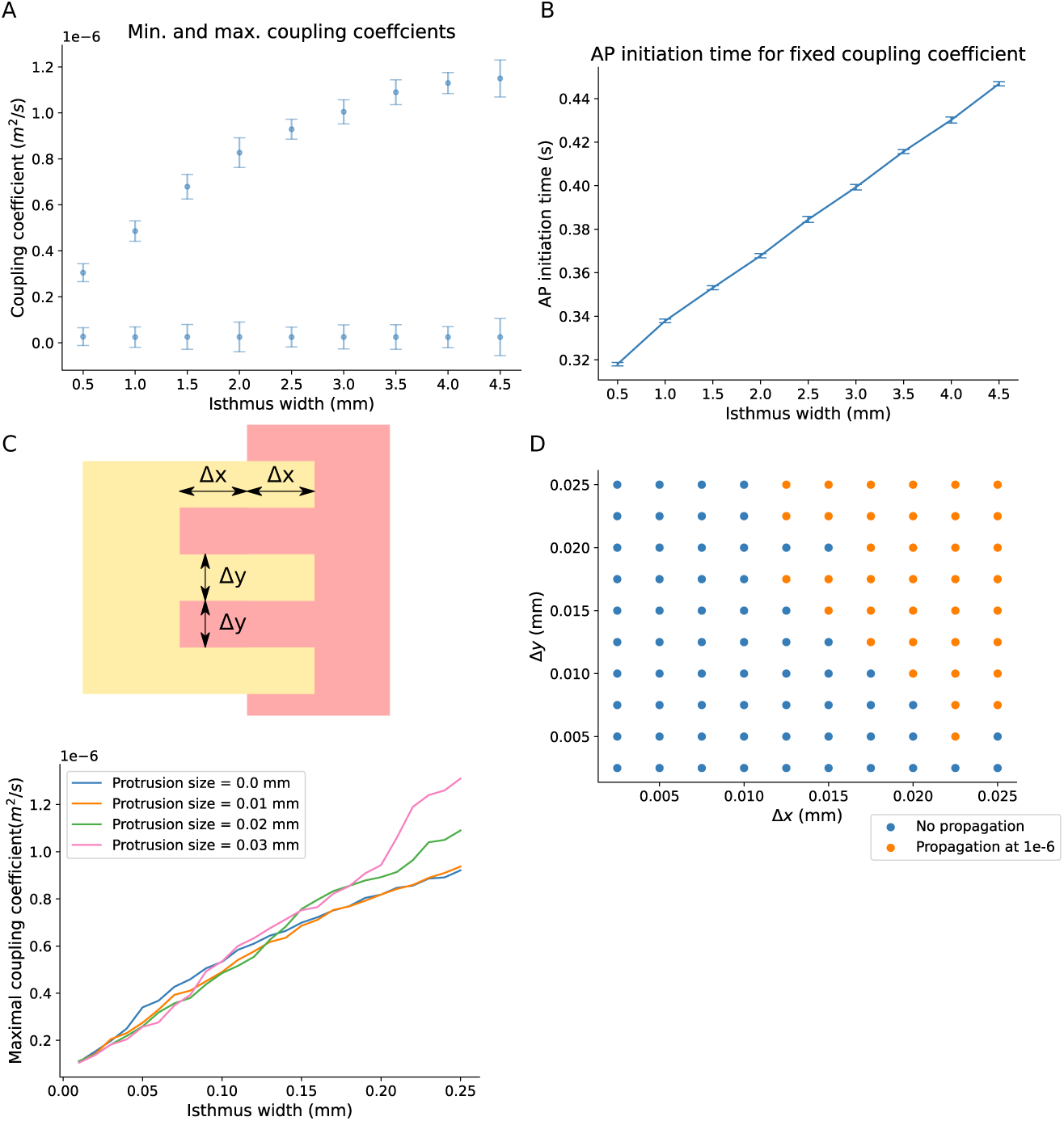
Action potentials propagate robustly across pacemaker-atrium boundaries with a broad isthmus. A: The minimal and maximal coupling coefficients for increasing isthmus width. All parameters were simulated 10 times for different random seeds. The electrophysiology was started after 5000 MCS. The bars indicate the standard deviation among these different simulations. B: The depolarization time for atrial cells with a varying isthmus width. C: Sketch of regular protrusion pattern, zoomed in around the end of the isthmus. The protrusions inward and outward have the same size Δx by Δy. D: Propagation or exit block plotted for different protrusion sizes. E: The maximal coupling coefficient that could be achieved with a fixed boundary between the pacemaker and atrial cells. Regular boundaries of different protrusion sizes were considered, indicated by the different colors.

**Fig 4.**
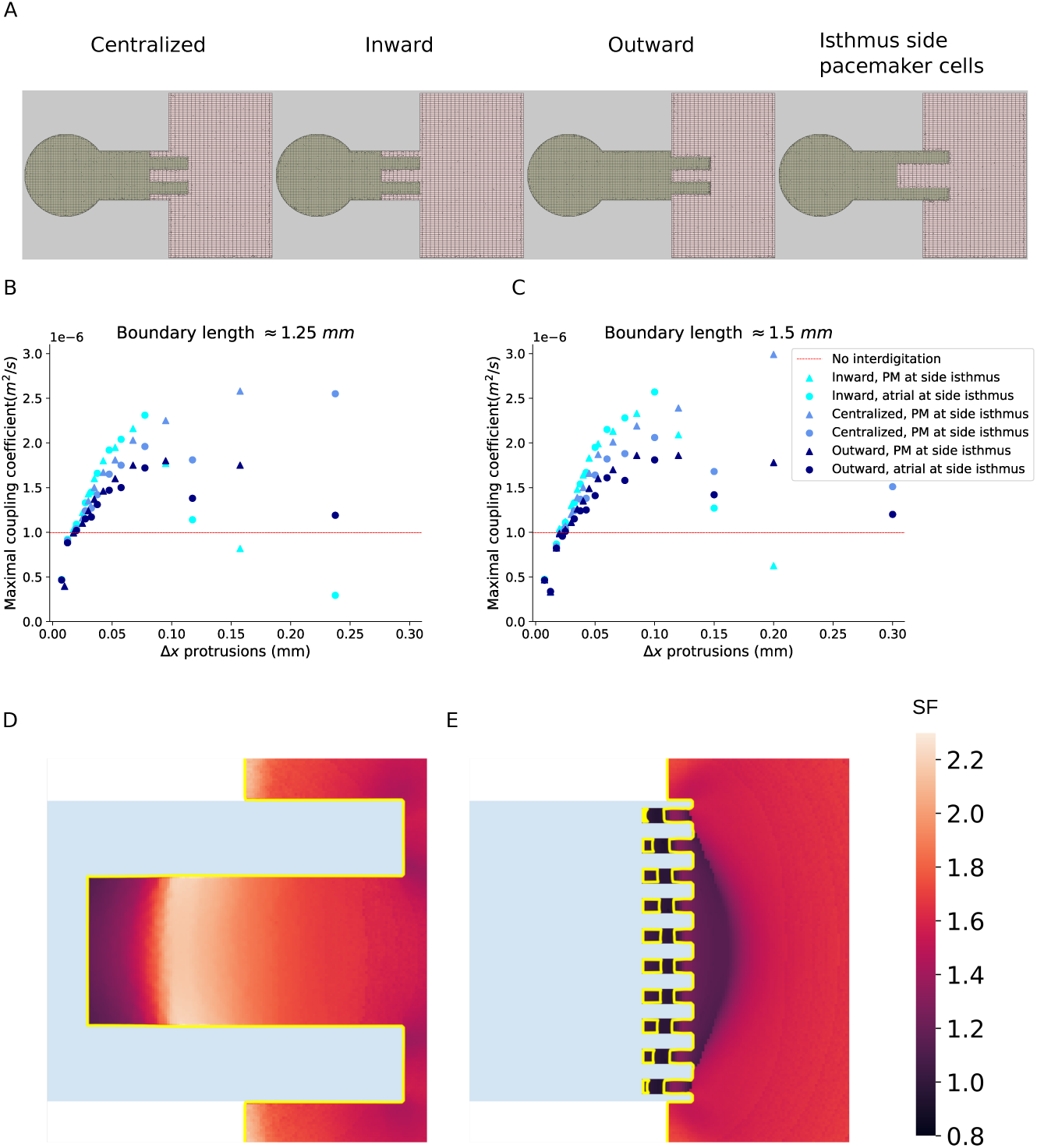
Medium-sized protrusions improve the robustness of propagation toward atrial cells. A: Examples of the ‘Centralized’, ‘Inward’, and ‘Outward’ configurations. B, C: The maximal atrial coupling coefficient such that an action potential still propagated for fixed total boundary length but protrusions of different sizes. The dashed red line indicates the mean of the maximal coupling coefficient and the standard deviation 10 repeats. B: For a total boundary length of 1.25 mm, C: For a total boundary length of 1.5 mm. The largest protrusions are not shown, because no successful propagation was found. D,E: Safety factor plots for large and small protrusions respectively. The yellow lines indicate the locations with SF = 1.

**Fig 5.**
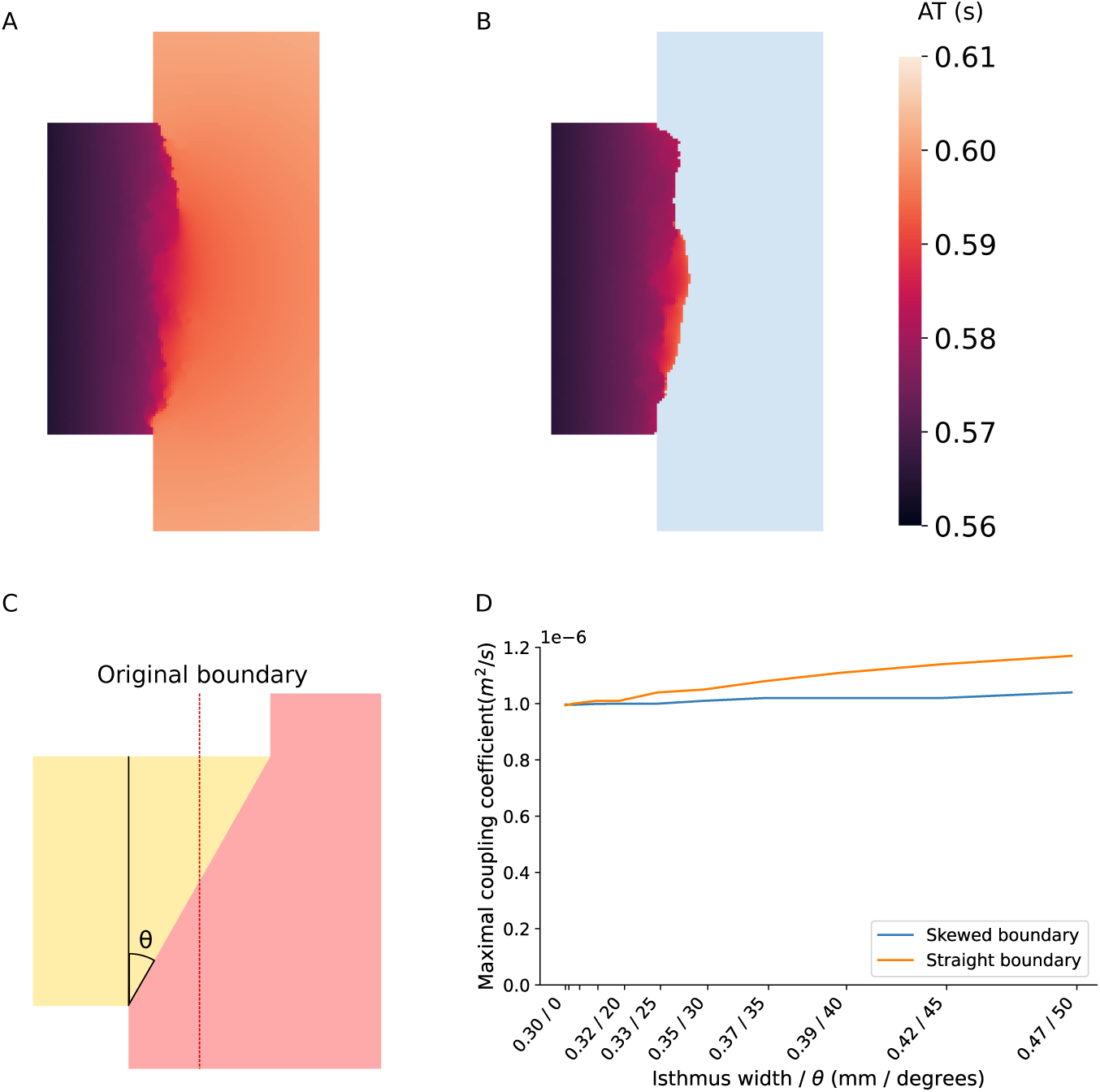
Action potentials propagate less robustly on boundary regions that activate asynchronously. A, B: Activation maps showing the elapsed time before a voltage of –40 mV is achieved for failed and successful propagation respectively on configurations with diffuse boundaries show asynchronous activation of cells near the end of the isthmus. C: Sketch of the skewed boundary configuration. The boundary was skewed with an angle θ. The number of atrial cells was kept constant by adjusting the shape of the atrial tissue. D: Comparison between the maximal coupling coefficients for skewed boundaries and straight boundaries of the same length indicates more robust propagation if the boundary length is increased with and without skewed boundaries. Action potentials propagated more robustly without skewed boundaries.

## Mathematical model

### Geometrical configuration

The geometry of the tissue represented a single SACP and consisted of a circular monolayer of pacemaker cells, a connecting isthmus of pacemaker cells, and a rectangular monolayer of atrial cells (Fig 1A). The geometrical configurations of the cells was initiated by random cell motility from the Cellular Potts model (CPM), a.k.a. Graner-Glazier-Hogeweg (GGH)-model [34, 35]. The CPM was combined with an electrophysiological model based on partial-differential equations following the work by Kudryashova et al. [36, 37] who introduced this type of model set-up to model cardiac monolayers and study propagation in highly fibrotic tissues. The CPM is a lattice-based model that represents cells are as patches of usually connected lattice sites. In the standard CPM, cells are subject to adhesive energy and an area constraint for which we used a standard Hamiltonian implementation [34, 35]. Furthermore, atrial cells are elongated, for which we added a length constraint [38] to the Hamiltonian:

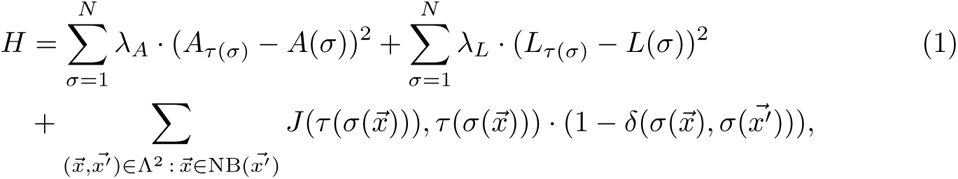

The amount of random cell motility in the CPM can be tweaked the parameter *T*, which determines the probability of accepting energetically unfavorable transitions. A list of CPM parameters is displayed in Table 1. After the initialization of the cellular configuration, the CPM is frozen and electrophysiology is started. In this work, we used the CPM to generate biologically-plausible SAN-atrium interfaces, after which we froze the CPM configuration and only ran the electophysiological model. Therefore, in this work we did not attempt to find realistic CPM parameters for pacemaker cells and atrial cells.

**Table 1.**
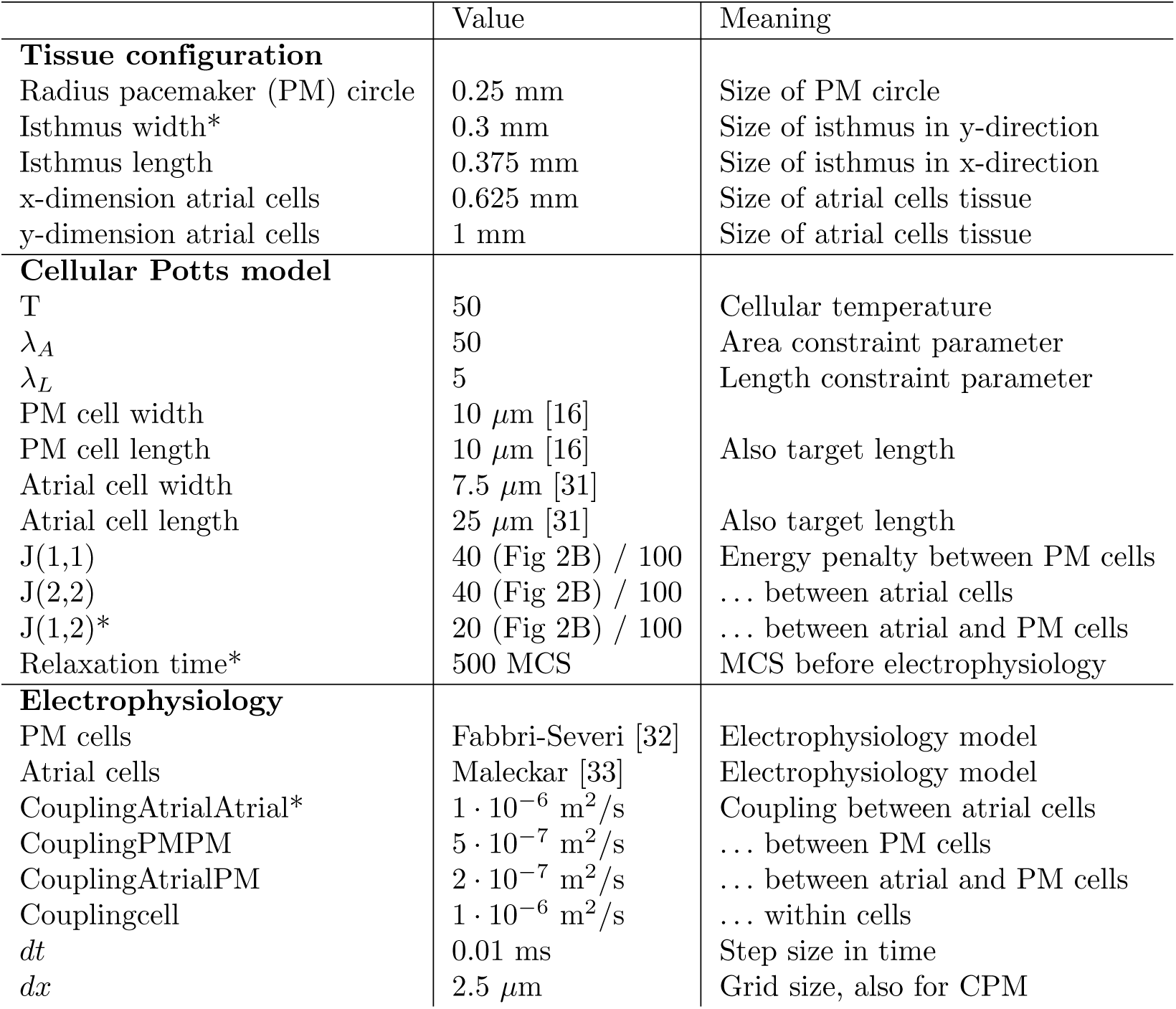
Standard parameters. Parameters with a * indicate that this parameter was varied in this manuscript; all other parameters were fixed throughout.

### Electrophysiology

Pacemaker cell electrophysiology was represented by the autocycling Fabbri-Severi model [32] and the atrial cell electrohypsiology was represented by the atrial cell model by Maleckar et al. [33]. Both of these models are monodomain models, i.e., only the intercellular voltages are computed and the extracellular voltages are not modeled explicitly. The models consist of a system of ordinary differential equations (ODEs) to describe the ionic currents, following the Hodgkin-Huxley formalism. The Fabbri-Severi model uses 33 variables and 11 ion currents to describe the electrophysiology, whereas the Maleckar model utilizes 30 variables and 13 ion currents. Both models hence provide a detailed description of the electrophysiology within these cells. To represent propagation of action potentials, the ODEs were converted into set of partial-differential equations (PDEs) using an operator splitting approach: The ODEs were evaluated inside each lattice point and these lattice points were coupled by allowing for a spatially varying diffusion of voltage between pixels with coupling coefficient D(⃗x). This results in the following set of partial differential equations (PDEs):

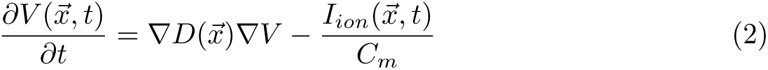

where *C_m_* is the membrane capacity. The first term in the derivative describes the change in voltage due to diffusion, while the second term describes the electrophysiology within this pixel. There is no need for an external activation current because the Fabbri-Severi model is autocycling. The coupling coefficient D(⃗x) is a property of a pixel that depends on the cellular configuration and differs between intracellular pixels and pixels at cell-cell interfaces. We distinguished between the following coupling coefficients, see (Fig 1B).

- Couplingcell: The coupling coefficient within cell interiors.
- CouplingAtrialPM: The coupling coefficient at the cell boundary between atrial and pacemaker cells.
- CouplingPMPM: The coupling coefficient in pixels at the cell boundary between pacemaker cells.
- CouplingAtrialAtrial: The coupling coefficient in pixels at the cell boundary between atrial cells.

The conduction velocity within the atrial tissue was largely determined by CouplingAtrialAtrial. A higher conduction velocity, however, also meant that ions diffuse rapidly to adjacent cells, making it more difficult to reach the initial activation threshold required to drive the atrial tissue. A list of the standard parameters we used can be found in Table 1. The coupling coefficients were chosen to achieve conduction velocities in the order of magnitude of 1-10 cm/s, which is consistent with conduction velocities found within *in vitro* monolayers [39, 40].

### Standard parameters

Table 1 provides a list of standard parameters used throughout this manuscript.

### Data analysis

The safety factor (SF) [41–45] provides a local measure of the robustness of propagation of an action potential. Intuitively, we define the safety factor as the fraction of the net amount of charge a pixel receives from its neighbors divided by the minimum charge required for activation if the pixel would be isolated. Hence, a safety factor larger than 1 indicates successful propagation and a higher safety factor indicates that the incoming charge exceeds the minimum required charge to activate this pixel. A high safety factor thus implies that a pixel is robust to small perturbations, i.e., it would still activate if the incoming charge would be reduced. On the other hand, a safety factor smaller than 1 indicates failure in activation. Because pacemaker cells will always spontaneously activate in isolation, the safety factor can only be applied to atrial cells. There exist many definitions of the safety factor [41–45]. We have adapted the safety factor as defined by Boyle et al. [45].

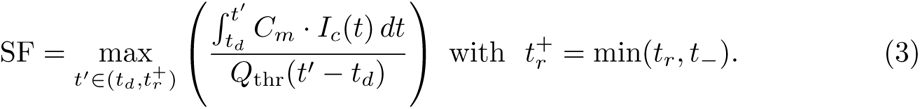

Here *t_d_* is the start of depolarization, defined as the time point when the voltage in a pixel crosses –70 mV from below. Similarly, *t_r_* is the repolarization time, defined as the time point when the voltage crosses –70 mV from above. *I_c_* indicates the membrane current due to diffusion through gap junctions, i.e., the net current entering a pixel at a time point due to diffusion. From this net current, we define t*_−_* as the moment in time when I_c_ becomes negative. Lastly, *Q_thr_*(*t^′^ − t_d_*) is the threshold charge required for the successful activation of a pixel. This threshold charge is determined by finding the minimal constant current such that applying this external current to an isolated pixel for *t^′^ − t_d_* time results in an action potential within 1 second of simulation time.

Although Eq 4 functions correctly for most pixels, the fraction that is maximized could have multiple local maximums for some pixels. For these pixels, the identified maximum before time t*_−_* may be a local maximum. For example, the global maximum could be achieved in the interval (*t_−_*, *t*^+^) if a small influx of charge is followed by large influx of charge. The pixels where a local maximum was found, were identified by a greatly reduced safety factor compared to its direct neighbors. For these pixels, the safety factors were recomputed with a computationally more expensive, method: First, a grid search is performed to sample the slope of the function in Eq 4, and thus estimate the locations of all local maximums on the interval (*t_d_*, *t_r_*). Then the exact values of all these local maximums are computed and safety factor is the global maximum among these local maximums.

Although the safety factor quantifies the local robustness of a propagating action potential, we are ultimately interested in the global robustness of an action potential as well. We assessed this by finding the largest sink that the pacemaker tissue could activate. The atrial tissue represents a voltage sink which is amplified if the cells are strongly electrophysiologically coupled. Better connectivity on the other hand results in higher conduction velocities, which is an essential property of the working myocardium. To assess how robustly an action potential propagates on a configuration, we identified the maximal coupling coefficient between atrial cells, couplingAtrialAtrial, that allowed for the propagation of the action potential (Fig 1C). This coupling coefficient describes the gap junctions between atrial cells, i.e., a large coupling coefficient corresponds to high gap junction expression. Action potentials dissipated in the atrial tissue if the coupling coefficient was too low because cells repolarized before transmitting the action potential to adjacent cells. On the other hand, exit blocks occurred for high coupling coefficients, because the atrial sink was too large in this case. If the coupling coefficient would be increased even further, the pacemaker cells would not activate spontaneously at all, because the hyperpolarization of the atrium would fully suppress the pacemaker cells. Only for intermediate values of the coupling coefficient, the pacemaker cells can successfully drive the atrial tissue. A higher value of this coupling coefficient indicates that the pacemaker cells could drive a larger sink and is hence more robust. This maximal value was found by initially taking a value that resulted in successful propagation and subsequently finding the maximal coupling coefficient that still resulted in successful propagation by a binary search.

### Implementation

We have used the Tissue Simulation Toolkit implementation [46, 47] of the Cellular Potts model for the initialization of our simulations. After this initialization, the electrophysiology was modeled as a layer on top of the resulting configuration. Instead of standard Monte Carlo sampling, we used the novel edge list algorithm [48] to improve computational efficiency. Furthermore, we implemented the local connectivity check by Durand and Guesnet [49] to prevent large atrial cells from engulfing small pacemaker cells and cell fragmentation due to the length constraint.

For the electrophysiological simulations, we used operator splitting in our computations. The ODEs were solved in parallel on a GPU using CUDA with the Forward Euler method. We took time steps of Δt = 10µs. The Rush-Larsen method [50] was also considered, but this decreased the stability of our simulations. Our spatial resolution was Δx = 2.5µm. The diffusion was computed with the Crank-Nicolson method [51]. This method requires inversion of tridiagonal matrices which was recently implemented efficiently in an interleaved format on CUDA [52]. The following procedure was repeated for evaluation of electrophysiological simulation.

1. Do a Forward Euler step of size 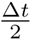 of electrophysiological activity.
2. Do the horizontal Crank-Nicolson step of size 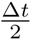.
3. Do a Forward Euler step of size 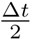 of electrophysiological activity.
4. Do the vertical Crank-Nicolson step of size 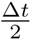.

This diffusion scheme is fully equivalent to the scheme used by Kudryashova et al. [36]. Because this entire procedure can be computed on the GPU, the communication between GPU and CPU is kept to a minimum, which further enhanced the simulation speed. Another advantage to the semi-implicit Crank-Nicolson scheme is that there is no hard limit on the maximal allowed time step size for stability such as the Courant number [53] for diffusion by forward Euler.

For the safety factor we had to compute *Q_thr_* every time we evaluated the value of the fraction in equation (3) because every pixel was in a different state. The cell state at the start of the safety factor measurement was taken as the initial state of the Maleckar model [33]. Consequently, we applied a constant stimulating current *I_stim_* (in pA/nF) for *t^′^ − t_d_* second of simulation time. If this resulted in successful activation of the pixel another attempt was made with a lower stimulating current. Alternatively, if the pixel did not get activated, another attempt was made with a higher stimulating current. With a binary search algorithm, the minimal activation current was found up to an accuracy of 10*^−^*^7^ pA / nF and we computed the threshold charge as *Q_thr_* = *I_stim_ ·* 1000 *· C_m_ ·* (*t^′^ − t_d_*). During a simulation, pixels do not receive a constant stimulating current from their neighbors like the test current supplied to compute the safety factor. Therefore, pixels with SF < 1 could still activate and pixels with SF > 1 may still fail to activate. The cut-off at SF = 1 should therefore be interpreted with caution. For pixels with several local maximums where the global maximum was not found with this method, we first did a grid search of evaluating the slope of the fraction in (Eq 4). This grid search procedure identified the approximate locations of local maximums. Consequently, the exact location of these maximums were again found by performing a binary search. The global maximum, and thus the safety factor, was identified as the largest value among these local maximums.

The simulation code and our implementation of the safety factor can be found on GitHub: https://github.com/Martijnadj/cardiac-interdigitation

## Results

### Action potentials on complex shapes due to random cell motility propagate less robustly

We first studied the model on diffuse pacemaker cell-atrium cell interfaces due to cell mixing, and found that diffuse interfaces had a negative effect on action potential propagation. All cells were initialized as rectangles and a random interface was generated from the CPM. For every simulation, a different interface was simulated. For some configurations, the atrial cells were excited whereas for others an exit block occurred (Fig 2A). The local robustness of action potential propagation at the end of the isthmus was quantified by computing the safety factor. As explained in detail in Section Data analysis, the safety factor is defined as the fraction between the charge delivered to a pixel and the minimal required charge required for activation. The yellow line shows the locations where the safety factor is 1, which indicates that the charge received in those pixels was almost, or just barely sufficient for the activation of this pixel. We see that in the case of successful activation, the safety factor is lowest near the isthmus. For the failed propagation, on the other hand, we see that the safety factor exceeds 1 only near the pacemaker. The source-sink mismatch was too large to propagate the action potential any further. Small differences in the boundary between the cell types can thus determine whether or not an action potential will propagate.

This was studied further by considering several surface tension parameters in the Cellular Potts model. The surface tension determines how strongly atrial and pacemaker cells will mix, and therefore how diffuse the tissue interface will become. We characterized the resulting interfaces by their total boundary length. The boundary length was defined as the total length of the interface between the two tissues, excluding the interfaces from dispersed cells. As expected, smaller surface tensions resulted in larger boundary lengths (S1 Fig). Electrophysiological simulations indicated that conduction blocks occurred more frequently for configurations with large boundary lengths (Fig 2B). We argued before that stronger coupling between atrial cells increases the strength of the current sink and thus complicates activation of the atrial tissue, see Materials and Methods for details. For some configurations with smaller boundary lengths, action potentials still propagated with a larger coupling coefficient between atrial cells, which further indicates that action potentials tend to propagate more robustly on configurations with smaller boundary lengths. Thus, contrary to our hypothesis, interdigitation may not necessarily benefit robust propogation of action potentials over the pacemaker-atrium interface. In the next Sections, we will study three properties of the interface geometry separately: (1) interfacial length; (2) interface morphology; (3) interface regularity. Firstly, interfacial length in itself may affect the robustness of action potentials. We will therefore study action potential propagation as a function of interface length, by varying isthmus widths while keeping interfacial boundary patterns constant. Secondly, with invariant interfacial length, the morphology of the interface may affect action potential propogation. We therefore studied action potential propagation for varying interfacial interdigitation patterns, while ensuring an invariant interfacial length. Thirdly, cell mixing may generate irregular interfaces, in which the action potential does not arrive at the interface everywhere synchronously. The effect of such asynchronous arrival of action potentials at the interface was studied in simulations with skewed boundaries

### Action potentials propagate more robustly for increased boundary lengths

After establishing that diffuse atrium-pacemaker boundaries reduce the robustness of action potential propagation, we next attempted to analyze (1) the effect of interface length, (2) the effect of interface morphology and (3) the effect of interface regularity. First we asked how the length of the pacemaker-atrium interface affects the robustness of action potentials by varying the width of the isthmus while keeping the morphology invariant. We either allowed the same random cell motility as in (Fig 1) or fixed a regular boundary as sketched in (Fig 3C). The robustness of the action potentials was studied by determining the strongest sink that the pacemaker could activate by increasing the connectivity between atrial cells. The strongest sink was found by performing a binary search as described in detail in Section Data analysis.

We found that a wider isthmus allowed for higher connectivity between atrial cells (Fig 3A) than a more narrow isthmus. The atrium gets activated normally (S1 Video) if we consider the maximal coupling coefficient, while an exit block occurs if the coupling coefficient were to be increased further (S2 Video). On the other hand, the lowest possible conduction coefficient that did not dissipate action potentials is independent of this isthmus width; this minimal coupling coefficient is a property of the atrial tissue instead. For this lowest possible coupling coefficient, the action potential can barely propagate through the atrial tissue (S3 Video), while for an even lower coupling coefficient, the traveling wave dissipates (S4 Video). Although a wide isthmus allows the pacemaker cells to drive strong atrial sinks, these sinks are also strongly connected to the pacemaker tissue because of the large surface between the cell types. Therefore, the hyperpolarization effect of the atrial cells inhibits the spontaneous activation of the pacemaker cells as can be seen in (S1 Video). For wider isthmuses, the depolarization time of pacemaker cells increased when using the maximal coupling coefficient for every isthmus. However, to confirm that this increased depolarization is not necessarily due to the stronger coupling between atrial cells, we also compared depolarization times for different isthmus widths with a coupling coefficient of 2 *·* 10*^−^*^7^. Indeed, we found that the wider connection between the tissues increases the depolarization times of the pacemaker cells and thus inhibits spontaneous activation (Fig 3B).

After establishing that wider isthmuses allow stronger electrical coupling and therefore faster propagation of action potentials from the pacemaker to the atrium, we next studied the role of the morphology of the interface. We varied the morphology of the interface by considering interdigitation patterns with digits of varying length and width. Contrary to the previous simulations, we explicitly fixed the boundary between the cell types, while still allowing random cell motility during initialization. Interdigitating patterns were constructed as shown in Fig 3C), where Δx represents the length of pacemaker tissue ‘digits’ into the atrium, and Δy represents digit width. Fig 3D) shows the results of a parameter sweep over Δx and Δy: this parameter sweep suggests that for a fixed isthmus width if the protrusions are long enough in the x direction, action potential propagation was possible. We have also tested protrusions of different sizes for a varying boundary length, where the protrusions had the same value of Δx and Δy. Similar to (Fig 3A), we found that a wider isthmus again allowed for larger coupling coefficients (Fig 3E). Additionally, larger protrusion sizes, where we fixed protrusion aspect ratio Δx = Δy, resulted in even better robustness for wider isthmuses.

### Action potentials propagate most robustly with medium-sized tissue protrusions

After establishing that longer interfaces allow stronger coupling between atrial cells, but inhibit spontaneous pacemaker activation, we next set out to study the effect of interfacial morphology in isolation from interface length and interface irregularity. To this end, we constructed a range of interdigitation patterns with varying Δx and Δy whilst keeping the total interface length constant, and again identified the maximum electrical coupling between atrial cells that still allowed for AP initiation in the atrial cells (Fig 4). Also we tested if the location of the interface within the isthmus affected the initiation of APs (Fig 4A). In the ‘centralized’ pattern, the middle of the patterns is located at the end of the isthmus. This ensured that all the patterns we tested had the same number of pacemaker cells. However, for larger values of Δx the pacemaker cells penetrated further into the atrial tissue than for smaller values of Δx, which may affect the results. As a control, we therefore also tested ‘inward’ patterns, in which pacemaker ‘digits’ terminate at the end of the isthmus and ‘outward’ patterns in which the base of the ‘digits’ is located the end of the isthmus (Fig 4A). Furthermore, the sides of this isthmus were either occupied by pacemaker or atrial cells to test all options with a fixed boundary length. Action potentials propagated less robustly if the sides were occupied by atrial cells (Fig 4B and 4C).

We tested AP initiation on interdigitation patterns ranging from the largest possible single protrusion up to protrusions of a single pixel size and maintained the total boundary length at 1.25 mm, compared to a length of 0.3 mm without digits, and again identified the maximum coupling coefficient that still allowed for robust initiation of APs in the atrial tissues. This boundary length choice resulted in atrial protrusions of the same order of magnitude as observed by Ten Velde et al. [23] for the medium-sized protrusions. Interestingly, such medium-sized protrusions of Δx between 0.0775 mm and 0.1575 mm allowed the optimal coupling coefficient (Fig 4B), whereas for smaller and larger protrusions the maximum coupling coefficient was lower. Furthermore, action potentials propagated more robustly on configurations where the isthmus sides were occupied by pacemaker cells rather than by atrial cells because the atrial cells were better surrounded by atrial cells for such configurations. Among the three interface variant, the ‘centralized’ interdigitated pattern allowed the strongest coupling, as indicated by the largest maximal coupling coefficient, with the maximal coupling coefficient for a straight boundary shown by the dashed red line for reference. The smallest protrusions result in less robust action potential waves than without any protrusions, while more robust action potential waves are found for tissues with medium-sized and large protrusion sizes.

We next asked if this observation depends on boundary length. (Fig 4C) shows the analysis for a larger boundary length of 1.5 mm. Larger boundary lengths than 1.5 mm physically did not fit in the configuration for the ‘inward’ and ‘outward’ patterns for the larges protrusions. Furthermore, in this case no propagating action potentials were found for the largest protrusion size of the ‘inward’ variant for any coupling coefficient, because the source-sink mismatch became too strong. Furthermore, for the largest protrusion, we found that the ‘centralized’ variant was also limited by the inhibition of spontaneous activity rather than exit blocks. That is, for these configurations, the ‘No activation’ regime as indicated in (Fig 1B) fully overlapped with the ‘Exit block’ regime. These results can be interpreted more easily by considering the safety factor. The safety factor was plotted in the vicinity of the end of the isthmus for large and small protrusion sizes. (Fig 4D and 4E) show safety factor patterns for the maximal coupling coefficient in both of these configurations. For the large protrusions, the local curvature effect caused a large surplus in charge within the atrial protrusion (Fig 4D). For the small protrusion size, the safety factor was less than 1 within the protrusions (Fig 4E). The hyperpolarizing effect of the pacemaker cells dominated the curvature effect of the protrusion, causing this configuration to propagate action potentials less robustly.

### Asynchronous action potentials propagate less robustly

Due to the irregular boundaries observed in (Fig 2), the boundary region between the cell types tended to activate asynchronously. Two examples, with an exit block and successful activation respectively, show that lattice sites near the end of the isthmus get activated asynchronously (Fig 5A and 5B). These figures display the first time a voltage of at least –40 mV is achieved. We hypothesized that these asynchronous bursts of action potentials were weaker than one well-synchronized front. Hence, the effect of asynchronicity was assessed by considering boundaries that were skewed by an angle θ. The number of atrial cells was kept constant by adjusting the shape of the atrial tissue (Fig 5C). Action potentials propagated more robustly if the the skewness of the boundary was increased. However, configurations with high skewness also have a larger boundary lengths. Since (Fig 3E) showed that increased boundary length improved robustness for regular boundaries, the effect of skewness should be compared with straight boundaries of the same length. This comparison suggests that although action potentials propagated more robustly with skewed boundaries than with straight boundaries, action potentials propagated even more robustly with straight boundaries with the same length (Fig 5D). This observation indicates that the increase in robust propagation was caused by an increase in boundary length while the asynchronicity due to skewness reduced the robustness of action potential propagation.

## Discussion

We have presented a hybrid model to study how interdigitation between SAN and atrium tissues could affect propagation of an action potential from a tissue consisting of pacemaker cells to a tissue consisting of atrial cells. This robustness was assessed locally by the safety factor and globally by finding the maximal intercellular coupling between atrial cells that allowed for propagation. Over diffuse tissue boundaries formed by random cell motility, action potentials (APs) tended to propagate more robustly over shorter tissue interfaces than over longer tissue interfaces. However, action potentials propagated more robustly through wide isthmuses (with longer tissue interface) than through narrow isthmuses with similar diffuse boundaries. We therefore concluded that apart from the length of the interface between pacemaker cells and the interface, also the interface morphology affected action potential propagation. To better understand the effect of interface morphology, we systematically varied interdigitation pattern morphology, while keeping the boundary length constant. We found that medium-sized protrusions improved the robustness of AP propagation, whereas small or large protrusions were less effective. Finally, we have shown that APs propagated less robustly if the isthmus boundary was activated asynchronously compared to straight boundaries of the same length.

### Effect of SAN-atrium interface morphology

The results from (Fig 2B) showed that action potential tended to propagate less robustly on tissues with long boundary lengths, which seems to contradict the hypothesis that interdigitation patterns improve how robustly action potentials propagate. We found two plausible explanations for the less effective AP propagation in these diffuse interface morphologies. Firstly, the protrusions formed by diffuse cell motility might be too small for effective AP propagation: (Fig 4) shows that APs propagate more robustly on configurations with a few large protrusions than on configurations with many small protrusions. Secondly, the diffuse morphologies had highly irregular protrusions of different sizes, with dispersed cells that had moved into the tissue of the other cell type. In these irregular configurations APs often arrived asynchronously at the atrial tissue. In (Fig 5) we have shown that asynchronous arrival of an action potential can reduce its robustness to propagate toward the atrium. This observation could explain the exit blocks in tissues with long but highly irregular boundaries between the cell types.

We also noticed that in (Fig 2) pacemaker cells diffused away from the main pacemaker tissue in some configurations. We hypothesized that such dispersed cells could hinder the propagation of action potentials. Such islands of pacemaker cells only had a noticeable effect on the robustness of action potential propagation if the islands were close together, effectively forming a barrier (S2 Fig). No significant differences were found if the barriers extended further vertically.

Apart from protrusion size, the location of the protrusion within isthmus affected AP propagation. All variants displayed optimal AP propagation for medium-sized protrusions (Fig 4B and 4C). For smaller protrusions, action potentials on the the ‘inward’ variant propagated more robustly with small protrusions than with the ‘centralized’ and ‘outward’ variants. With large protrusions on the other hand, action potentials on the ‘inward’ variant propagated less robustly than on the ‘centralized’ and ‘outward’ variants. This difference may be explained by the fact that protrusions were better insulated from the atrial tissue in the ‘inward’ variant compared to the other variants. On the other hand, the atrial cells were located closer to the pacemaker cells for large protrusions, which caused the atrial cells to inhibit spontaneous pacemaker activity if the intercellular connection was too strong. For a boundary length of 1.5 mm, no action potentials initiated for the largest protrusions on the ‘inward’ because the atrial cells suppressed spontaneous pacemaker activation. We also found that action potentials on the ‘centralized’ variant propagated more robustly than on the ‘outward’ variant for the same protrusion size. This difference was expected since the protrusions in the ‘outward’ variant were less insulated from the atrial cells than in the ‘centralized’ variant. This last observation underlines the importance of the location of interdigitation patterns within sinoatrial conduction pathways. If interdigitation patterns were located at the end of the sinoatrial conduction pathway, action potentials would propagate less robustly than when the interdigitation patterns were located inside the insulated pathway. For *in vitro* studies, the best performing location of the interdigitation pattern could therefore depend on the size of the protrusions. For small protrusions, our simulations predict that embedding the interdigitations within the isthmus gives the most robust AP propagation, whereas for larger protrusions, more robust AP propagation is predicted if these protrusions are located at the end of the isthmus. This difference may be explained by the suppressing effect of atrial cells close to the SAN center for large protrusions.

### Related work

Although our geometrically detailed *in silico* study of the sinoatrial conduction pathways is novel, many other simulation studies of the SAN have been performed before (reviewed in Ref. [54]). In particular, Winslow and Varghese already studied the effect of interdigitation [28]. To study interdigitation, they compared an *in silico* SAN with and without atrial protrusions. They found that the SAN with atrial protrusion activated the surrounding atrium more effectively when these atrial protrusions were included. Huang and Cui performed a more extensive parameter study [29]. For different coupling strengths between cells, they identified ‘no pacing’, ‘pacing and driving’, ‘pacing and no driving’ parameter regimes. Upon adding 1-dimensional strands from the atrium towards the SAN center, they found that the ‘pacing and no driving’ parameter regime increased whereas the ‘no pacing’ regime increased. Our study supports both of these previous *in silico* studies on interdigitation. Furthermore, we claimed that interdigitations were effective because pacemaker cells surrounded the atrial strands. This argument is supported by [55], who simulated a tree graph of atrial and pacemaker cells to study the effect of simultaneous activation of atrial cells. They found that if the first atrial cells were surrounded by more atrial cells, activation of atrial cells was more robust.

However, these studies did not consider the effect of the sinoatrial conduction pathways. The malicious and beneficial effects of SACPs have been studied at length. Kharche et al. studied how a combination of multiple active SACPs could lead to micro-reentry [13]. An accurate reconstruction of the SAN was activated externally. Because of the interaction between the SACPs, micro-reentry around the SAN was observed. Zyantekorov et al. studied the effect of SACP width on action potential propagation to the atrium [14]. Wide SACPs suppressed spontaneous pacemaker activation more strongly than narrow SACPs, while wide SACPs simultaneously improved the activation of the atrium. The balancing between these two regimes highlights the importance of a the micro-structure within SACPs. A heterogeneous fibrotic SACP was studied by Li et al. [12]. They studied the effect of reduced sodium currents and increased adenosine concentrations with and without fibrosis. They found that fibrotic SANs were more vulnerable to these changes, which is consistent with the *in vivo* study they performed. The boundary region between the pacemaker and atrial cells was studied more comprehensively by Amsaleg et al. [15]. The gradient and mosaic model were tested in an 3D SAN which was connected to the atrium with 5 SANs. They succeeded to achieve activation of the crista terminalis with one specific combination of sigmoidal gradients. Possibly, the 3D configuration necessitated a well-tuned gradient. More likely, however, is that the structure within the SAN is more complex in reality.

One such way to add complexity is to add cellular detail to electrophysiological simulations such as performed by Kudryshova et al. [36, 37]. The gap junctions in cardiomyocytes are non-uniformly distributed along the cell membrane. Kudryshova et al. implemented this anisotropicity in gap junctions [36] by modeling cardiac monolayer tissues with a Cellular Potts model [34, 35]. They constructed monolayers of cells which matched experimental monolayers of cardiac cells. These *in silico* monolayers were later adapted to study the propagation of action potentials in monolayers with a high percentage of fibroblasts [37]. By including a mechanism for cytoskeleton alignment, they achieved conduction with higher percentages of fibroblasts than predicted by previous model. This underlines the importance of a sufficient level of detail to fully understand the propagation of action potentials. We hence followed this approach as we were interested in relatively small protrusions to study interdigitation. On the other hand, our configuration of SACPs was inspired by previous *in vitro* [30] and *in silico* [12] studies. This combination of cellular and electrophysiological detail on the one hand, but *in vitro*-inspired configurations on the other hand enabled us to suggest design principles for *in vitro* monolayer studies.

### In vitro and in vivo validation

We studied interdigitation in isolation of other effects by excluding transitional cell types, contrary to Winslow and Varghese [28]. With this approach we tested the interdigitation hypothesis and found that larger boundary lengths are beneficial. Moreover, we could conclude that protrusions sizes as observed *in vivo* [23] are more robust than smaller or larger protrusions. Lastly, our fine spatial scale approach enabled us to observe asynchronous action potential fronts and consequently we found that well-synchronized action potentials indeed improve robustness of AP propagation. These insights how geometry affect the robustness of action potentials could only be achieved with a detailed geometrical approach. A 1D model such as the model by Huang and Cui [29] would miss such effects. A 3D model would possibly discover even more geometrical effects which cannot be found with our 2D model.

Despite the increased robustness of AP propagation over an interdigitated SAN-atrium interface, interdigitation is not necessary to drive a tissue of atrial cells. Amsaleg et al. [15] studied electrophysiological models in a more realistic 3D geometry than ours with a straight, diffuse interface and still managed to drive their atrial tissue. Instead of interdigitation, the authors achieved successful AP propagation into the atrium by implementing both a gradient in coupling coefficients and gradient in the prevalence of both cell types around the boundary of the two cell types. Our results from (Fig 2C) suggest that a mosaic pattern may hinder AP propagation, which may explain why Amsaleg et al. did not find full AP propagation for other sigmoidal gradients. Their approach of a gradient models may be more realistic than our approach because some aspects of transitional cell types have been observed in human hearts [18, 22]. Transitional cells should therefore be included in future interdigitation models when building a more realistic model of the human SAN.

Although transitional cell types have been observed *in vivo*, explicit transitional cell types are not strictly necessary to achieve a gradient in electrophysiological properties. In fact, the boundary region between two cell types can display properties intermediary to both cell types by exchanging ions. These implicit intermediary cells are demonstrated in a study by Ori et al. [56]. In this study, kidney cells are studied which cannot propagate an action potential on their own. They modified these kidney cells by either adding a sodium or potassium current. Cells with both currents could propagate an action potential, while cells with only one of these currents could not. Yet, at the interface of those two cell types an action potential could propagate, which indicates that sufficient ions are exchanged to effectively obtain transitional cells. Furthermore, the AP morphology at the interface between pacemaker and atrial also changes gradually from a pacemaker action potential to an atrial action potential as found *in vivo* [21, 23, 57] and also simulated by *in silico* studies [10, 12, 15, 29, 58]. Additionally, the average cell in the boundary region has a lower coupling coefficient than atrial cells, but higher than pacemaker cells. We therefore argue that interdigitation is an alternative method to achieve a smooth transition region between pacemaker cells and atrial cells, rather than an explicitly gradient in coupling coefficients and electrophysiology. This further explains why medium-sized protrusions could propagate action potentials most robustly as more ‘effectively transitional’ cells are present with medium protrusion sizes compared to other protrusion sizes.

## Limitations of the study

### Safety factor

The safety factor was used to identify the local robustness of AP propagation. It also highlights the curvature effect in large protrusion. However, the method has limitations. Firstly, intuitively the value SF = 1 should be a sharp cut-off between activation and non-activation. However, (Fig 4E) indicates that this is not always the case. The pixels within the protrusion got activated in this simulation, but have a safety factor smaller than 1. This counterintuitive result is caused by the method *Q_thr_* is determined. This charge is computed by only considering constant stimulating currents. However, if a pixel would receive a short burst of external current, it may still be activated without receiving the total charge predicted by *Q_thr_* because the total charge leaking by depolarizing currents would be lower than in the case of a constant current. Secondly, we did not succeed in finding a catch-all solution for our maximization problem for the safety factor while keeping the algorithm computationally feasible. This high computational cost is the result of our choice to compute *Q_thr_* separately for every pixel. An alternative method to reduce computational cost would be to sample *Q_thr_* for many initial values of the ODE model and many different stimulation times. For an unknown value of the ODE variables and stimulation times, the *Q_thr_* could then be interpolated instead of requiring expensive computations. However, this method would only be worthwhile when sampling the safety factor many times. Furthermore, the linear interpolation of unknown parameters may be inaccurate because the system is sensitive to small variable changes. A more sophisticated machine-learning approach could perhaps predict the safety factor better. Lastly, for pixels where a simple binary search did not find the global maximum, we suggested an alternative method for safety factor computation by first performing a grid search. This method required small step sizes, and thus long computation times, for some pixels because I_c_ can oscillate in inhomogeneous grids. However, despite these computational problems, the safety factor remains a useful tool identify on which part of the atrial tissue action potentials propagate most robustly.

### Low conduction velocities

Although in the human heart typically conduction velocities of 40-90 cm/s (right free atrium) are found [39, 59–61] and 100-120 cm/s in the crista terminalis [59, 61]. [39, 59–61], we could only study the effect of relatively small coupling coefficients compared to other *in silico* studies [14, 15, 36]. This in turn resulted in lower conduction velocities of the action potential (typically in the range of 1-10 cm/s). A key difference between the *in vivo* heart and our study is that the human heart is three-dimensional, while we have modeled a monolayer of cells. In monolayer *in vitro* cultures, however, the conduction was found to be variable among different human stem-cell derived cardiomyocytes: 7.1 *±* 2.4 cm/s [39], 8.8 *±* 1 cm/s [39], 17.64 *±* 0.89 cm/s [40]. Hence, although our conduction velocities were much lower than those found *in vivo*, they were of the same order of magnitude as *in vitro* monolayer cultures. We hypothesize that we could not achieve propagating action potentials for high conduction velocities partly because the model was two-dimensional. Higher conduction velocities required higher coupling coefficients which either resulted in an exit block or hyperpolarization of the pacemaker (Fig 1B). Three-dimensional models on the other hand tend to allow for higher conduction velocities [13, 15]. Furthermore, we opted to ignore potential gradients in coupling coefficients, while there are clues that such gradients exist and could be relevant.

### Future prospects

The current model could be extended to investigate interdigitation combined with other possible explanations such as transitional cell types, or extended to three dimensions to eventually mimic the complex three-dimensional nature of the *in vivo* sinoatrial node. Also, we have paid limited attention to the creation of interdigitation patterns. Random cell motility from the Cellular Potts model trivially resulted in some interdigitation if we tuned our adhesion parameters to allow mixing between pacemaker cells and atrial cells. However, observed interdigitation patterns are unlikely to be caused by this mechanism, as the tissues would continue to mix more in this case while cardiac tissues are immobile and the protrusions were relatively small. Further study on the formation of interdigitation patterns would be required to better understand the role of such patterns. Another possible topic for future research would be to further identify the best types of patterns. We limited ourselves to a small number of configurations we thought to be relevant while many more configurations for the boundary patterns are possible. The full range of possibilities could for instance be systematically explored with machine learning techniques to find the most robust boundary patterns.

## Conclusion

The goal of this paper was two-fold: on the one hand, we aimed to understand the effect of interdigitation on the propagation of action potentials from pacemaker cells to atrial cells. On the other hand we chose to stick closely to *in vitro* monolayer cell cultures to suggest design principles that could inform such studies. The global robustness of the action potential on a configuration was assessed by determining the maximal coupling coefficient that still allowed the action potential to propagate. We have shown that, although the total boundary length increased, diffuse boundaries due to random cell motility decreased the robustness of action potential propagation. Consequently, we have isolated the effect of the boundary length by increasing the width of the isthmus and found that this improved the robustness instead. Furthermore, when we isolated the effect of interdigitation, we found that action potentials propagated most robustly on configuration with medium-sized protrusions. Small protrusions with the same boundary length did not necessarily improve the robustness of the action potential, and neither did the largest protrusions. We also found yet another explanation for the decreased propagation of action potentials on configurations with diffuse boundary regions. In these configurations, action potentials arrived at the boundary between the cell types asynchronously and this asynchronicity reduced the robustness of action potential propagation. All in all, this indicates that action potentials propagate more robustly on configurations with interdigitation between pacemaker cells and atrial cells, but only if the digits are large enough and synchronicity is preserved. Future research could further identify how this principle could be of importance *in vitro* and *in vivo*, or could attempt to further elucidate the intricacies of interdigitation *in silico*.

## Supporting information

**S1 Fig.**
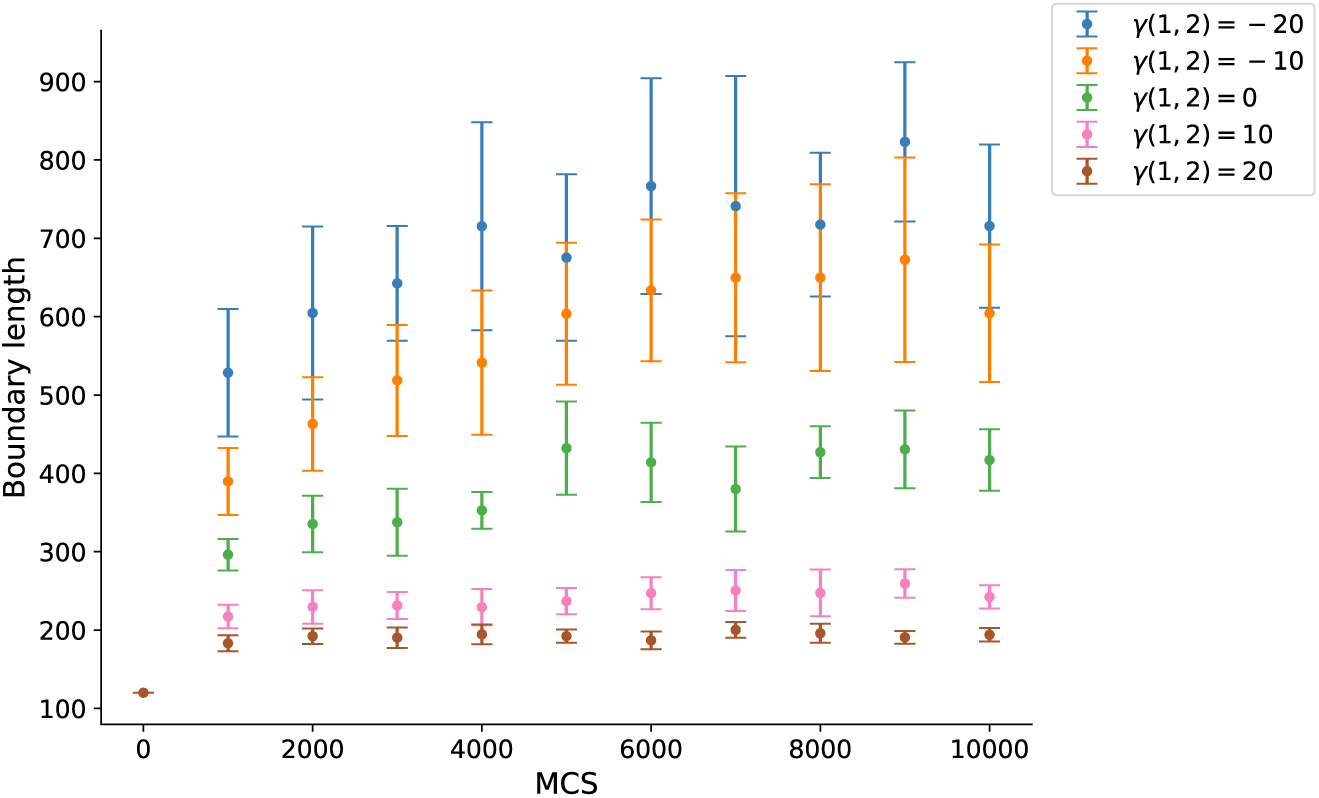
Larger boundary length and heterogeneous cellular interfaces reduce propagation robustness. The boundary length over time for different surface tensions (n = 10). Loose (clusters of) cells are excluded from this boundary length. For negative surface tensions, the boundary length increases over time, while the boundary length remains approximately constant for surface tensions larger or equal to 0.

**S2 Fig.**
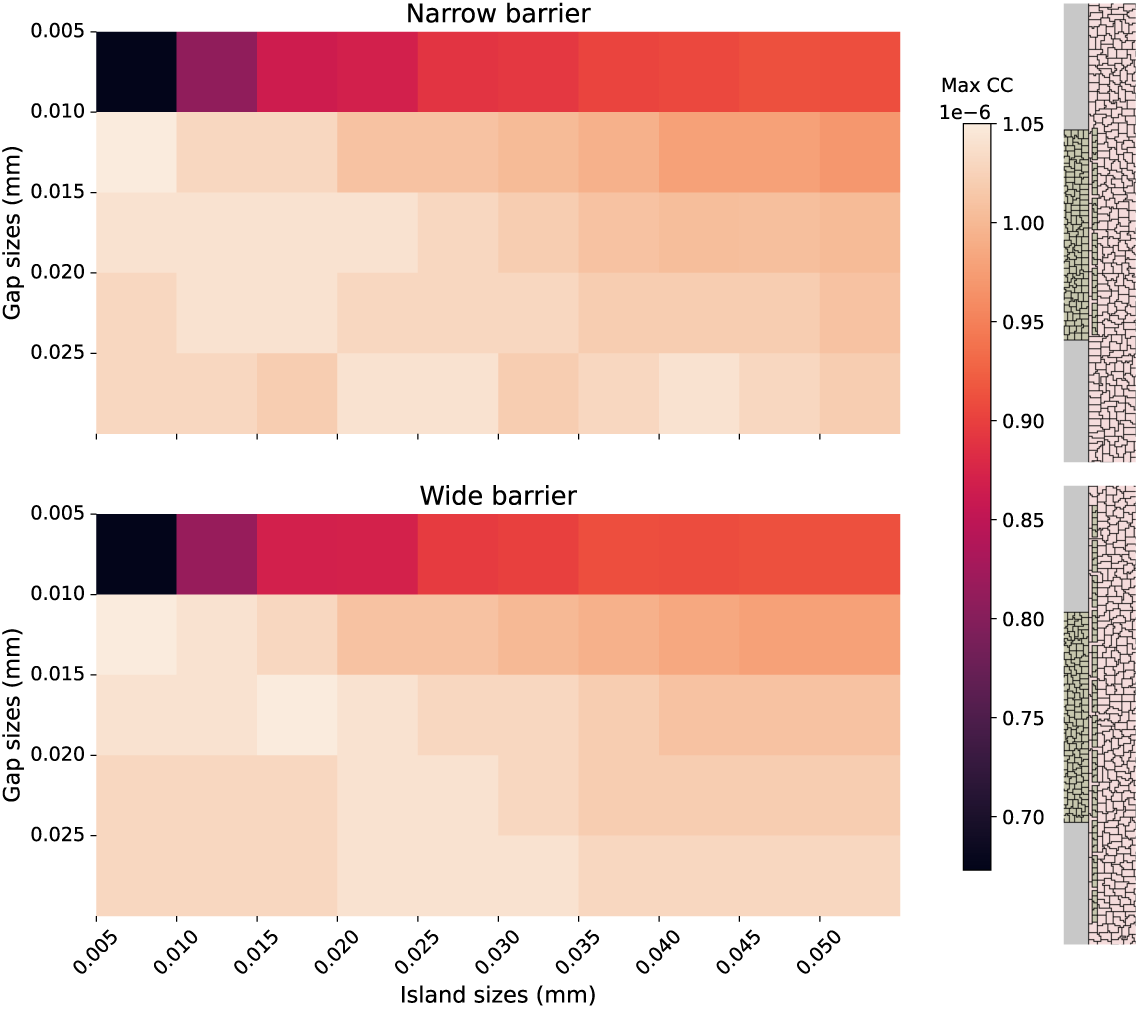
Maximal coupling coefficient such that action potentials propagate when islands of pacemaker cells are placed after the isthmus. Action potentials get hindered by clusters of pacemaker cells if the clusters are close to each other. No differences are observed when this barrier is extended past the width of the isthmus.

**S1 Video. Action potentials propagate normally with maximal coupling coefficients.** A different maximal coupling coefficient is found for every isthmus width. A wider isthmus allows for a higher maximal coupling coefficient and higher conduction velocities.

**S2 Video. Action potentials are blocked at the end of the isthmus with coupling coefficients larger than the maximal coupling coefficients.** For every isthmus width, a coupling coefficient slightly higher than the maximal coupling coefficient that would allow propagation is chosen. Exit blocks occur at these coupling coefficients.

**S3 Video. Action potentials with the lowest possible coupling coefficients.** The lowest coupling coefficient is similar for all isthmus widths. The action potentials propagate but are unstable.

**S4 Video. Action potentials with lower coupling coefficients than required for propagation.** The action potentials dissipated for these coupling coefficients. This lower bound is a property of the atrial tissue rather than the transitional area because similar minimal coupling coefficients are found for all isthmus widths.

## Acknowledgments

This work was performed using the compute resources from the Academic Leiden Interdisciplinary Cluster Environment (ALICE) provided by Leiden University.

This work was supported by NWO grant NWO/ENW-XL 2019.029 (MdJ and RMHM), and by Prof. dr. Jan van der Hoevenstichting voor Theoretische Biologie (RMHM) affiliated to the Leiden University Fund (RMHM).

