## Supplementary figures and images for "In silico model suggests that interdigitation promotes robust activation of atrial cells by pacemaker cells"

### Supplemental Figure 1

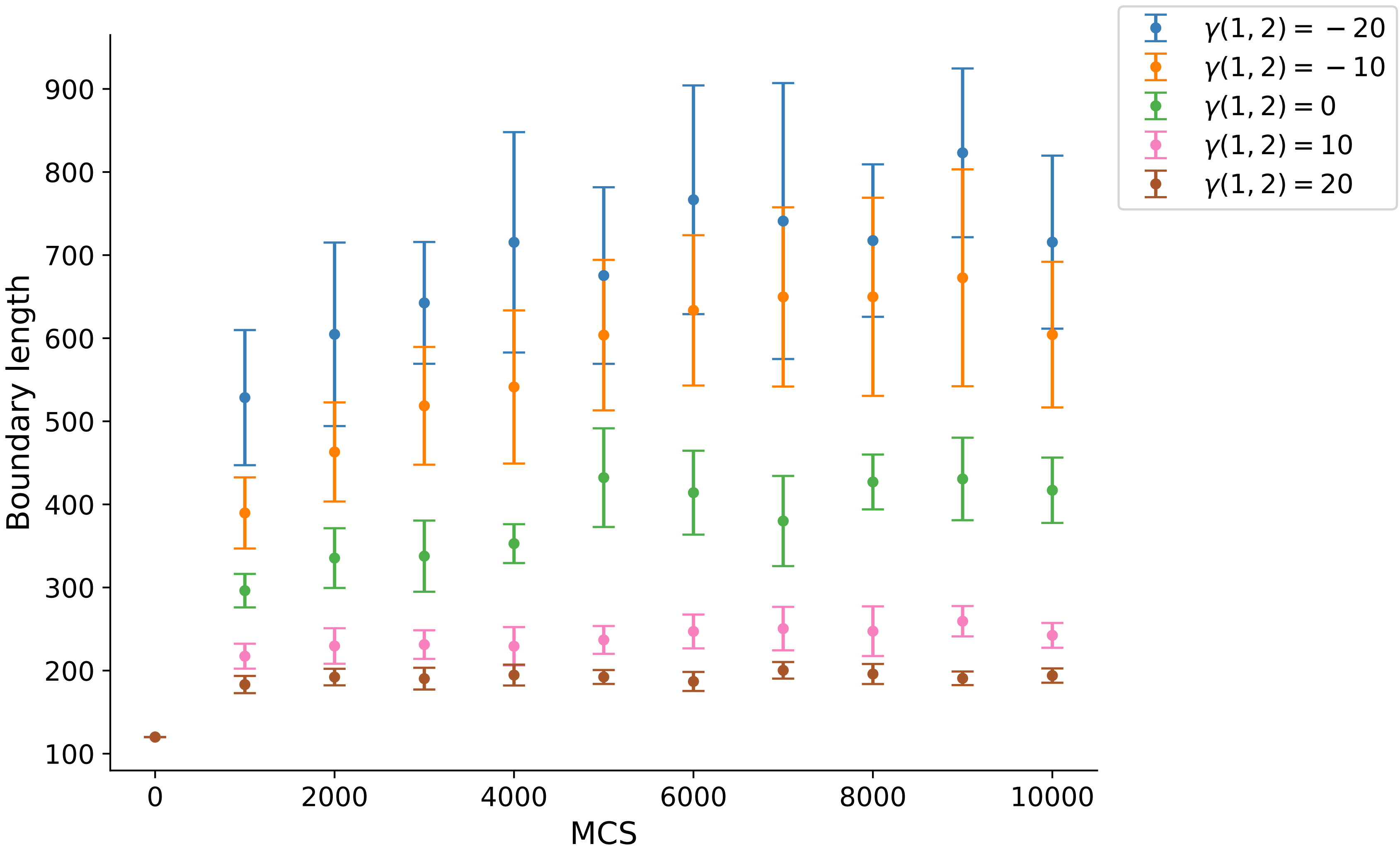

### Supplemental Figure 2

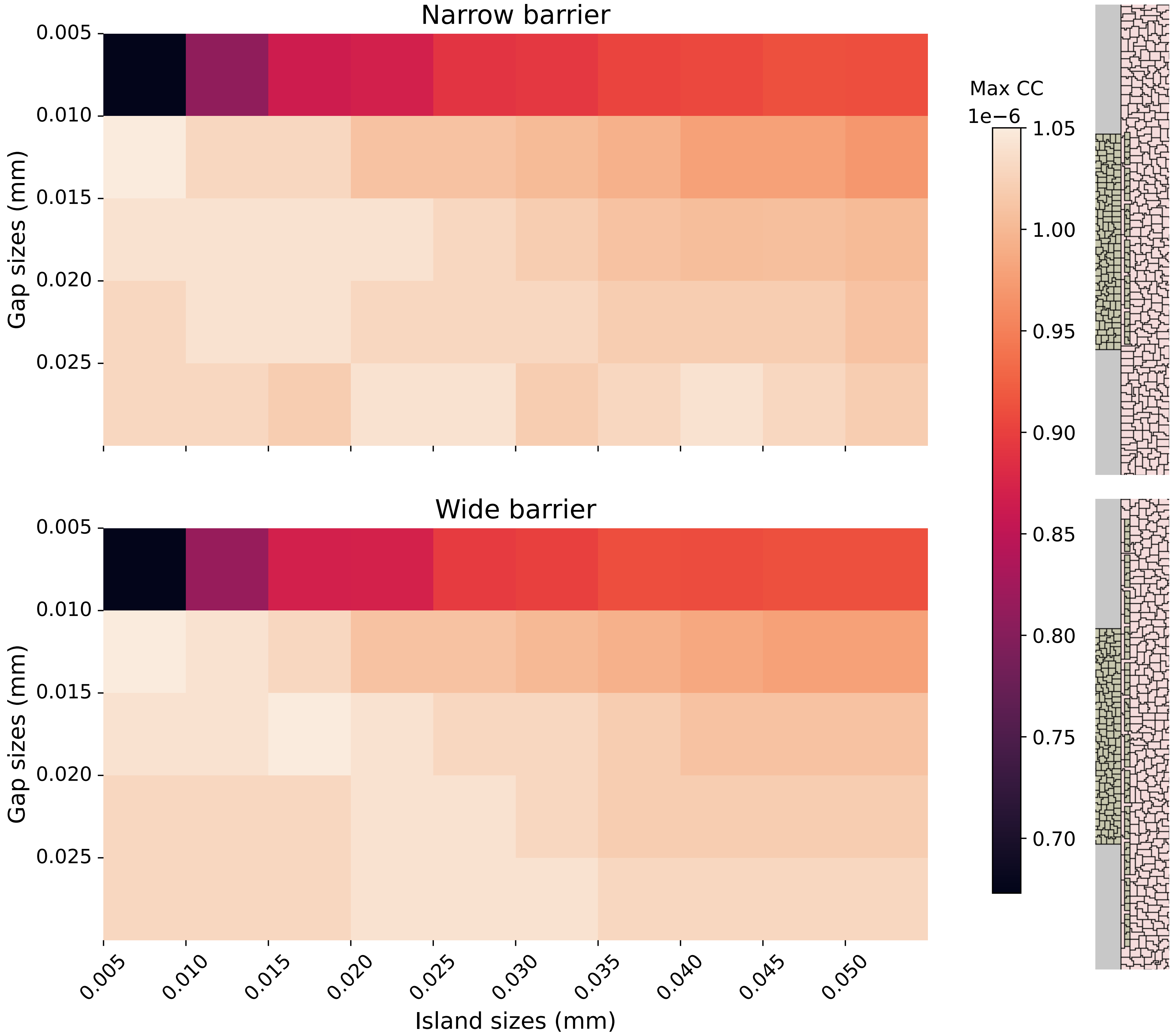
